# Exhaustive capture of biological variation in RNA-seq data through k-mer decomposition

**DOI:** 10.1101/122937

**Authors:** Jérôme Audoux, Nicolas Philippe, Rayan Chikhi, Mikaël Salson, Mélina Gallopin, Marc Gabriel, Jérémy Le Coz, Thérèse Commes, Daniel Gautheret

## Abstract

Each individual cell produces its own set of transcripts, which is the combined result of genetic variation, transcription regulation and post-transcriptional processing. Due to this combinatorial nature, obtaining the exhaustive set of full-length transcripts for a given species is a never-ending endeavor. Yet, each RNA deep sequencing experiment produces a variety of transcripts that depart from the reference transcriptome and should be properly identified. To address this challenge, we introduce a k-mer-based software protocol for capturing local RNA variation from a set of standard RNA-seq libraries, independently of a reference genome or transcriptome. Our software, called DE-kupl, analyzes k-mer contents and detects k-mers with differential abundance directly from the raw data files, prior to assembly or mapping. This enables to retrieve the virtually complete set of unannotated variation lying in an RNA-seq dataset. This variation is subsequently assigned to biological events such as differential lincRNAs, antisense RNAs, splice and polyadenylation variants, introns, expressed repeats, and SNV-harboring or exogenous RNA. We applied DE-kupl to public RNA-seq datasets, including an Epythelial-Mensenchymal Transition model and different human tissues. DE-kupl identified abundant novel events and showed excellent reproducibility when applied to independent deep sequencing experiments. DE-kupl is a new paradigm for analyzing differential RNA-seq data with no preconception on target events, which can also provide fresh insights into existing RNA-seq repositories.

## Background

Successive generations of RNA sequencing technologies have established since the 1990’s that organisms produce a highly diverse and adaptable set of RNA molecules. Modern transcript catalogs such as Gencode[1] now reach hundreds of thousands of transcripts, reflecting widespread pervasive transcription and alternative RNA processing. However, in spite of years of high throughput sequencing efforts and bioinformatics analysis, we contend that large areas of transcriptomic information remain essentially disregarded.

To illustrate this point, let us consider the biological events that drive transcript diversity. Firstly, transcripts result from transcription initiation events either at promoters of protein-coding and non-coding genes, or at multiple antisense or inter/intra genic loci. Secondly, transcripts are processed by a large variety of mechanisms, including splicing and polyadenylation, editing [2], circularization [3] and cleavage/degradation by various nucleases [4, 5]. Thirdly, an essential, yet often overlooked, source of transcript diversity, is genomic variation. Polymorphism and structural variations within transcribed regions produce RNAs with single nucleotide variations (SNVs), tandem duplications or deletions, transposon integrations, unstable microsatellites or fusion events. These events are major sources of transcript variation that can strongly impact RNA processing, transport and coding potential.

Current bioinformatics protocols for RNA-seq analysis do not properly account for this vast diversity of transcripts. Prevalent computing strategies can be roughly classified into two categories: reference-based tools [6, 7, 8, 9] rely on the alignment (or pseudo-alignment) of RNA-seq reads to a reference genome or transcriptome, while de novo assembly tools [10] reconstruct full-length transcripts based on the analysis of RNA-seq reads. These protocols fail to account for true transcriptional diversity in several respects: (i) they ignore small-scale variations such as SNP or indels (ii) they rely on full-length transcripts that cannot represent the full range of variation observed in organisms and (iii) they misrepresent transcripts containing repeats due to ambiguity in alignment or assembly.

We propose a new approach to RNA-seq analysis that facilitates the discovery of any type of event occurring in an RNA-seq library independently of alignment or transcript assembly. Our approach relies on k-mer indexing of sequence files, a technique that recently gained momentum in NGS data analysis [8, 9, 11, 12, 13]. To identify biologically meaningful transcript variation, our method selects k-mers with differential expression (DE) between two experimental conditions, hence its name: DE-kupl. Using public human RNA-seq datasets, we show that a large amount of RNA variation can be captured that is not represented in existing transcript catalogs. As proofs of concept, we applied DE-kupl to RNA-seq data from an Epythelial-Mensenchymal Transition (EMT) model and from different human tissues. DE-kupl identified abundant novel events and showed excellent reproducibility when applied to independent deep sequencing experiments.

## Results

### Reference datasets are an incomplete representation of actual transcriptomes

First, we analyzed k-mer diversity in different human references and high-throughput experimental sequences. To this aim, we extracted all 31-nt k-mers from sequence files using the Jellyfish program [14]. Figure 1A-B compares k-mers from Gencode transcripts, the human genome reference and RNA-seq libraries from 18 different individuals [15] corresponding to three primary tissues (6 libraries/tissue). To minimize the risk of including k-mers containing sequencing errors, we retained for each tissue only the set of k-mers that appear in 6 or more individuals.

**Figure 1.**
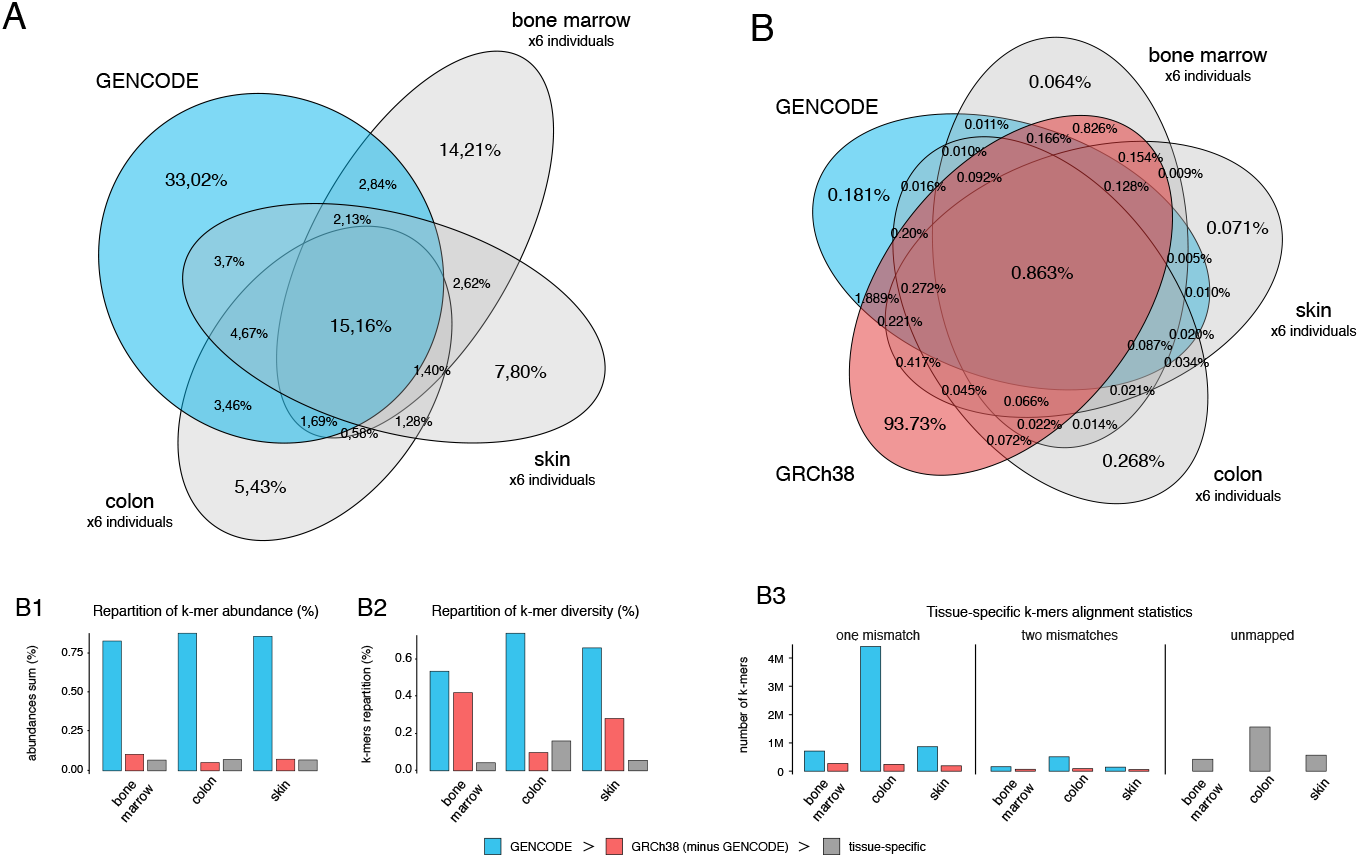
The diversity of 31-mers in RNA-Seq libraries exceeds that of reference sequences. **A**. Intersection of k-mers present in Gencode transcripts and RNA-Seq data from three tissues: bone marrow, skin and colon. The set of k-mers for each tissue was defined as the common k-mers shared by all six individuals. **B**. Intersection of k-mers present in Gencode transcripts, the reference human genome (GRCh38) and RNA-Seq data (same as in A). **B1**. Repartition of k-mer abundances for each tissue represented in A and B. K-mers shared with Gencode are labelled as “GENCODE”, among other k-mers, those shared with the human genome are labelled as “GRCh38”, the remaining k-mers are labelled as “tissue-specific”. The same procedure was applied in B2 and B3. **B2**. Repartition of k-mer diversity for each tissue. **B3**. Mapping statistics of k-mers labeled as “tissue-specific” in B2. These k-mers were first mapped to Gencode transcripts, and unmapped k-mers were then mapped to the GRCh38 reference using Bowtie1, allowing up to 2 mismatches in a 31-mer.

Measures of k-mer abundance show that k-mers are overwhelmingly associated to Gencode transcripts (Fig 1B1). However, when considering k-mer diversity, a large fraction of k-mers are tissue-specific and not found in the Gencode reference (Fig 1A). These tissue-specific k-mers may result from sequencing errors, genetic variation in individuals or novel, or non-reference transcripts. The majority of RNA-seq k-mers that do not occur in Gencode are found in the human genome reference (Fig 1B, 1B2), suggesting polymorphisms and errors represent a minor fraction of tissue-specific k-mers and a lot of k-mers results from expressed genome regions that are not represented in Gencode. Further scrutiny of tissue-specific k-mers shows that a significant fraction can be mapped to the transcriptome with one substitution. However, for each tissue there is an average of 1 million k-mers that cannot be mapped to either reference (1B3).

Non-reference k-mers classify samples as accurately as reference transcripts. We performed a Principal Component Analysis (PCA) of the above human tissue samples using conventional transcript counts and k-mer counts. PCA based on 20,000 randomly selected unmapped k-mers was able to differentiate tissues as well as PCA based on estimated gene expression or transcript expression (Fig 2). This illustrates how a “shadow”, non reference transcriptome that is not incorporated in standard analyses comprises biologically relevant expression data.

**Figure 2.**
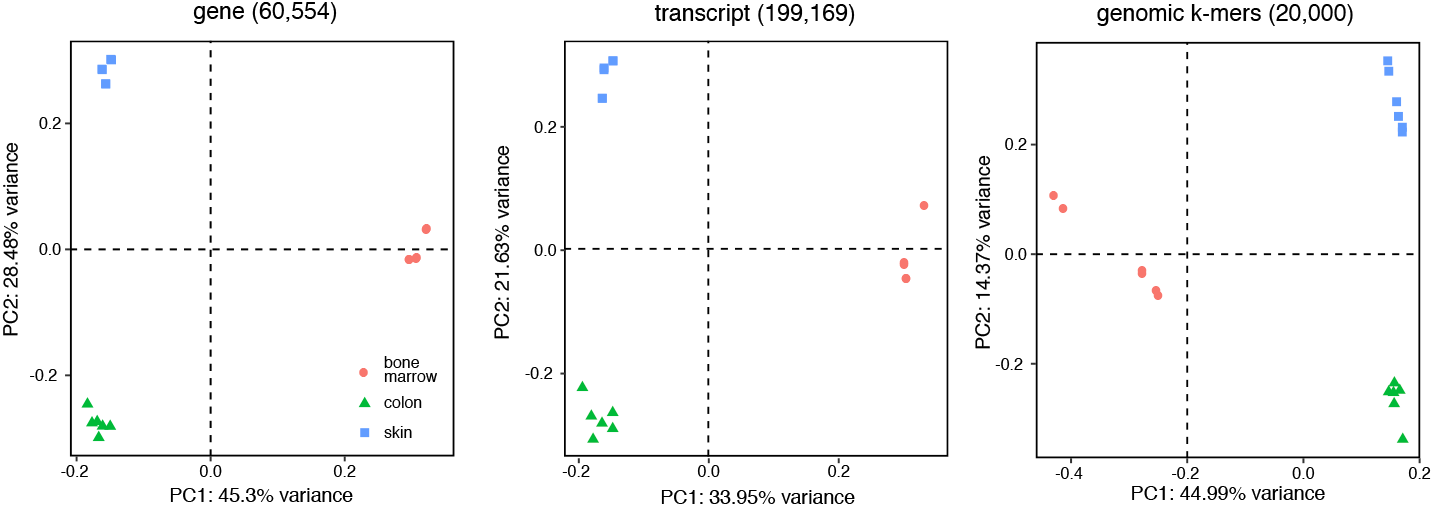
Principal Component Analysis on non-reference k-mers discriminates tissues. Samples are labeled according to their tissues (bone marrow, colon, skin). PCA were produced with normalized, log transformed counts. For genes and transcripts, counts were generated with Kallisto based on Gencode V25. Genomic k-mers correspond to 20k random k-mers from the RNA-seq libraries that did not map to Gencode transcripts but successfully mapped to GRCh38.

When comparing RNA-seq and whole genome sequence (WGS) data from the same individual [16], library-specific k-mers represent a much larger fraction of RNA-seq than of WGS k-mers (Fig 3). This shows that non-reference sequence diversity is larger in RNA-seq than in WGS. Altogether these results point towards the existence of a significant amount of untapped biological information in RNA-seq data.

**Figure 3.**
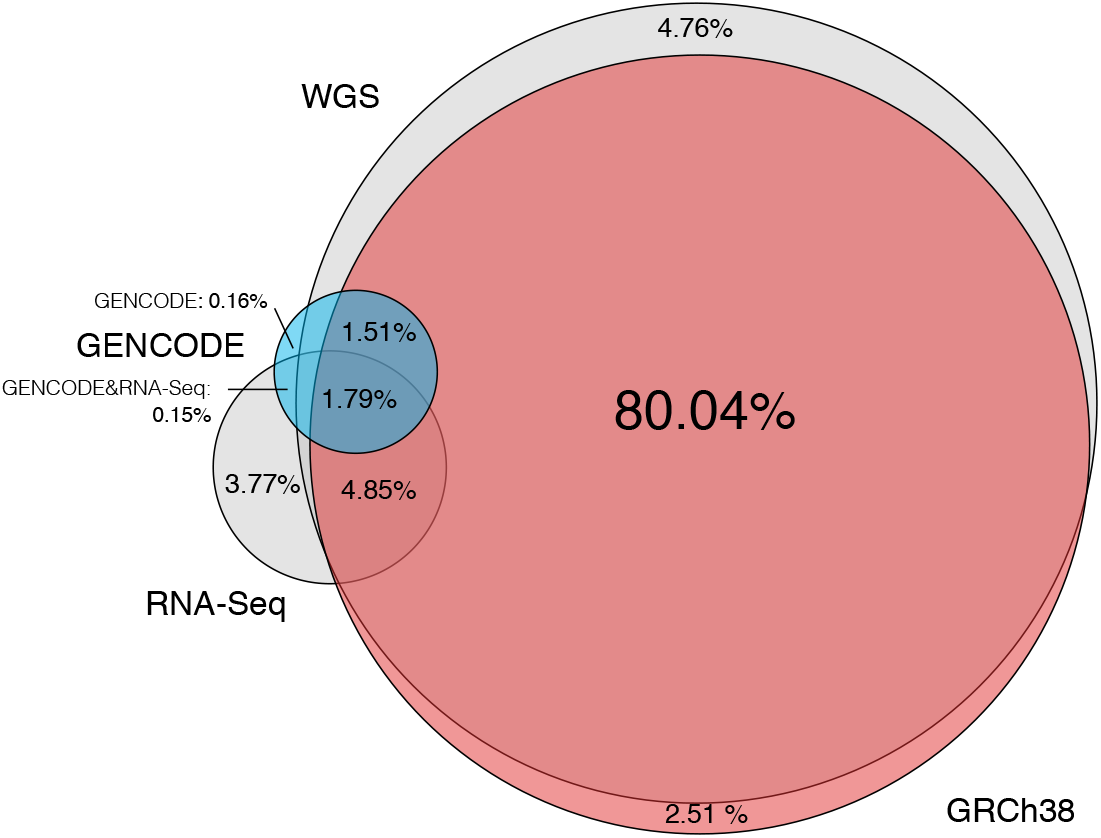
The diversity of non-reference k-mers is greater for RNA-Seq than for whole genome sequencing (WGS). Intersection of k-mers between Gencode transcripts, the human genome (GRCh38), RNA-Seq and WGS data. RNA-Seq and WGS data originate from the same lymphoblastoid cell line (HCC1395).

Non-reference k-mers may result from three classes of biological events. First, they may stem from genetic polymorphism in the studied sample. Second, they may result from RNA processing, notably, but not limited to, splicing and polyadenylation. A predominant source of k-mers in this category is intron retention, whose products are not usually incorporated into reference databases and are mostly by-products of regular gene expression. A third, major source of k-mer‘innovation” is intergenic expression (eg. lincRNA, antisense RNA, expressed repeats or endogenous viral sequences). Altogether, the combination of these genetic, transcriptional and post-transcriptional events may have a profound impact on transcript function.

### A new k-mer based protocol for deriving transcriptome variation from RNA-seq data

We designed the DE-kupl computational protocol with the aim to capture all k-mer variation in an input set of RNA-seq libraries. This protocol is composed of four main components (Figure 4):

1. Indexing: index and count all k-mers (k=31) in the input libraries
2. Filtering: delete k-mers representing potential sequencing errors or perfectly matching known transcripts
3. Differential Expression (DE): select k-mers with significantly different abundance across conditions
4. Assembly and annotation: build contigs of assembled k-mers and annotate contigs based on sequence alignment.

**Figure 4.**
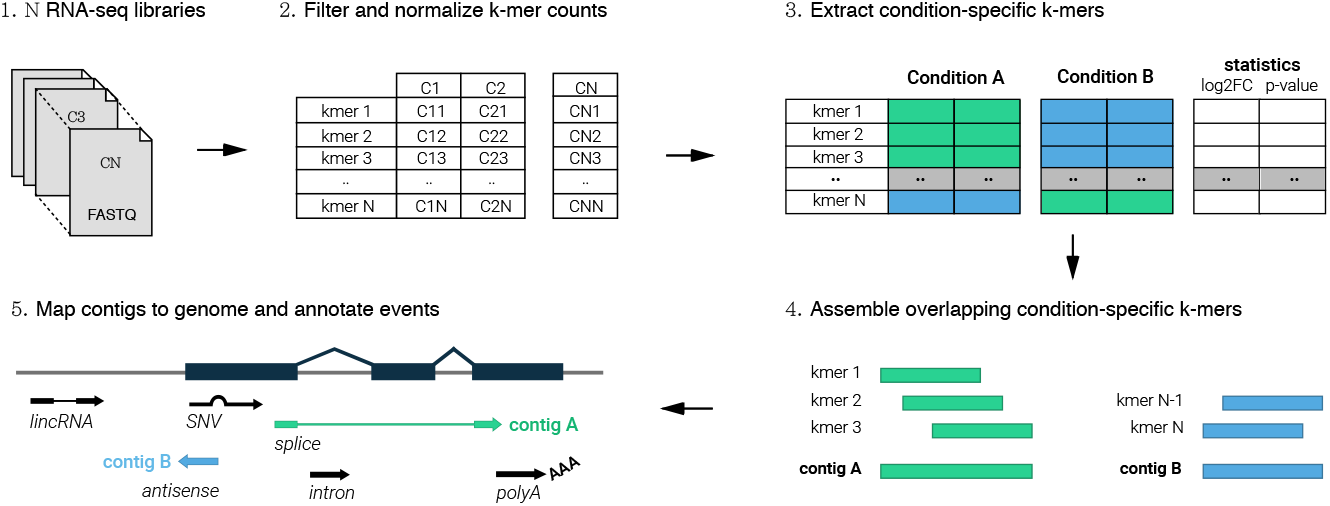
The DE-kupl pipeline for the discovery and analysis of differentially expressed k-mers. First, Jellyfish is applied to count k-mers in all libraries. K-mers counts are then joined into a count matrix and filtered for low-recurrence and matching to the reference transcriptome. Normalization factors are computed from raw K-mer counts and the DE procedure is applied. Finally overlapping DE k-mers are merged into contigs and annotated based on their alignment to reference and overlap with annotated genes

DE-kupl departs radically from existing RNA-seq analysis procedures in that it does neither “map-first” (a la Tuxedo suite [17]) or “assemble-first” (a la Trinity [18]) but instead directly analyzes contents of the raw FASTQ files, displacing assembly and mapping to the final stage of the procedure. In this way, DE-kupl guarantees that no variation in the input sequence (even at the level of a single nucleotide) is lost at the initial stage of the analysis. Even unmappable k-mers such as sequences from repeats, low complexity regions or exogenous organisms, are retained up to the final stage and can be analyzed. The DE-kupl protocol is detailed in Methods. We highlight here some of its key features. First, DE-kupl must deal with the large size of the k-mer index. A single human RNA-seq library contains in the order of 10^8^ distinct k-mers and an index for 50 individual samples can reach billions of k-mers. We selected the Jellyfish tool for counting k-mers [14] as it presents very fast computing times and allows to store the full index on disk for further query.

The central process in DE-kupl is k-mer filtering. Filtering out unique or rare k-mers is relatively straightforward and considerably reduces k-mer diversity and the amount of sequence errors. Another stringent filter is the removal of k-mers matching reference Gencode transcripts. The rationale for this is that the bulk of k-mers in RNA-seq data comes from expressed exons, and we are not interested in this canonical exon expression, as it can be captured efficiently by conventional, reference-based protocols [8, 9]. Discarding these k-mers enable us to ignore the overwhelming signal caused by known transcripts and focus on expressed regions harboring differences from the reference transcriptome.

Two modes are available to perform differential analysis of k-mers (Figure S10 & Methods): The *t*-test filter mode is fast and has lower sensitivity, i.e. it retrieves only the most significantly differentially expressed k-mers. The DESeq2-based mode [19] is slower, more sensitive and is recommended for small samples (fewer than 6 vs. 6 samples). Whenever possible, key steps of the procedure (k-mer table merging, t-test, k-mer assembly) were written in C, enabling the whole procedure to run on a relatively standard computer in a reasonable amount of time.

### Discovery and assembly of differential RNA contigs with DE-kupl

To assess the capacity of DE-kupl to discover novel differential events, we applied the procedure to 12 RNA-seq samples from an EMT cell-line model [20], in which NSCLC cells were induced by ZEB1 expression over a 7-day time course. We compared 6 RNA-seq libraries from the “Epythelial” stage of the time course (uninduced and Day 1) with 6 libraries from the “Mesenchymal” stage (Day 6 and 7). The full DE-Kupl procedure was completed in about 4 hours in the *t*-test mode (single-threaded), and 6.5 hours in the DESeq2 mode (multi-threaded), using 8 computing cores, 54 GB RAM and 7 to 42 GB of hard disk space (Table 1). Recurrence filters efficiently reduced k-mer counts from 707M to 92.5M and the Gencode filter further reduced counts to 40.3M. Differential analysis using the *t*-test mode eventually retained 3.8M k-mers that were assembled into 133.690 contigs (Table 2). The resulting contigs ranged in size from 31 bp (corresponding to an ‘orphan” k-mer) to 3.6 kbp, with a major peak of short 31-40 bp contigs and a minor peak around 61 bp (Fig 5A).

**Figure 5.**
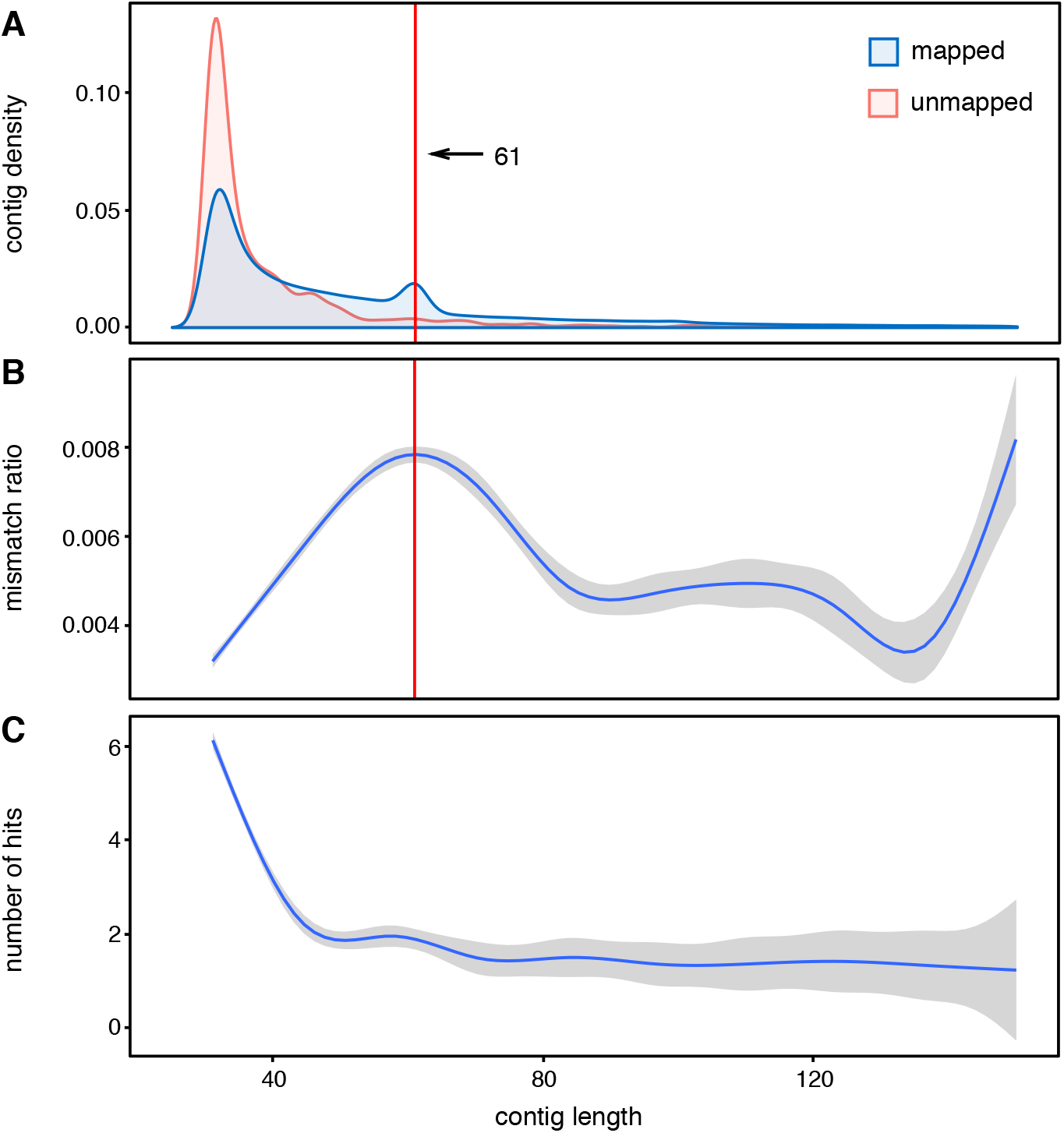
Specificity of differentially expressed contig. **A**. Density plot of contig lengths for mapped and unmapped contigs. The red line indicates contigs assembled from k k-mers and likely corresponding to SNVs. **B**. Mismatch ratio (number of mismatch / contig size) as a function of contig length. **C**. Number of hits in the reference genome as a function of contig length. The B and C curves were obtained using a smoothing function.

**Table 1.** DE-kupl parameters and resources used for analyzing EMT data (12 libraries) using the *t*-test and DESeq2 methods.

| Parameter/Resources | value |  |
| --- | --- | --- |
| nb_threads | 8 |  |
| min_recurrence | 6 |  |
| min_recurrence_abundance | 5 |  |
| pvalue_threshold | 0.05 |  |
| lib_type | stranded |  |
|  | *t*-test | DESeq2 |
| Max memory usage | 54Gb | 53Gb |
| Max disk used (1) | 7Gb | 42Gb |
| Running time (1) | 4h 02m | 6h 33m |

(1): excluding reference genome and transcriptome indexing for the annotation step.

**Table 2.** DE-kupl pipeline results for the EMT experiment. Description of output files sequentially generated by DE-kupl. Number of kmers/contigs, correspond to the number of lines in each file.

| Files | Description | Number of k-mers/contigs |  | Sizes |  |
| --- | --- | --- | --- | --- | --- |
| raw_counts (no filter) | Matrix of k-mers counts from all libraries | 707,067,278 |  | (not generated) |  |
| raw_counts.tsv.gz | Matrix of all k-mer counts from all libraries with recurrence filters | 92,525,450 |  | 1.9GB |  |
| noGENCODE-counts.tsv.gz | Counts filtered with GENCODE k-mers | 40,398,848 |  | 728MB |  |
|  |  | *t*-test | DESeq2 | *t*-test | DESeq2 |
| diff-counts.tsv.gz | Counts with differential expression test, filtered on adjusted P value | 3,813,418 | 6,102,447 | 186MB | 510MB |
| merged-diff-counts.tsv.gz | DE k-mers assembled into contigs | 133,690 | 169,613 | 3.0MB | 18MB |

Almost all (99.2%) of the 133k DE contigs mapped to the human genome. Mapping revealed that most 61 bp contigs result from assembly of 31 overlapping k-mers harboring a single nucleotide variation (SNV) at every position of the k-mer. This phenomenon also causes a higher mismatch ratio for contigs around 61 bp (Fig 5B). Contigs that do not map to the human genome are generally shorter than mapped contigs (Fig 5A), indicating a lower signal-to-noise ratio in unmapped contigs. Expectedly, shorter mapped contigs tend to map at multiple loci more often than longer ones (Fig 5C), however 80% of all contigs are uniquely mapped (not shown).

Analysis of contig locations reveals distinct contig classes. Most contigs are located in annotated introns and exons (Fig 6), however intronic contigs are predominantly exact matches while exonic contigs are predominantly mismatched. This effect is due to Gencode filtering: contigs with exact matches to introns are usually not filtered, as they do not pertain to a Gencode transcript, while contigs that match exons are filtered out unless they differ from the reference. This difference might be in the form of SNVs, or through exons extending in flanking intergenic or intronic regions. Under the same rationale, contigs mapping to intergenic and antisense regions are depleted in SNVs (Fig 6), consistent with their location in unannotated lncRNAs and antisense-RNAs, while contigs overlapping exon-exon junctions behave like exonic contigs (high rate of SNV). However, a significant fraction of exon junction contigs are exact matches, indicating they may correspond to novel junctions.

**Figure 6.**
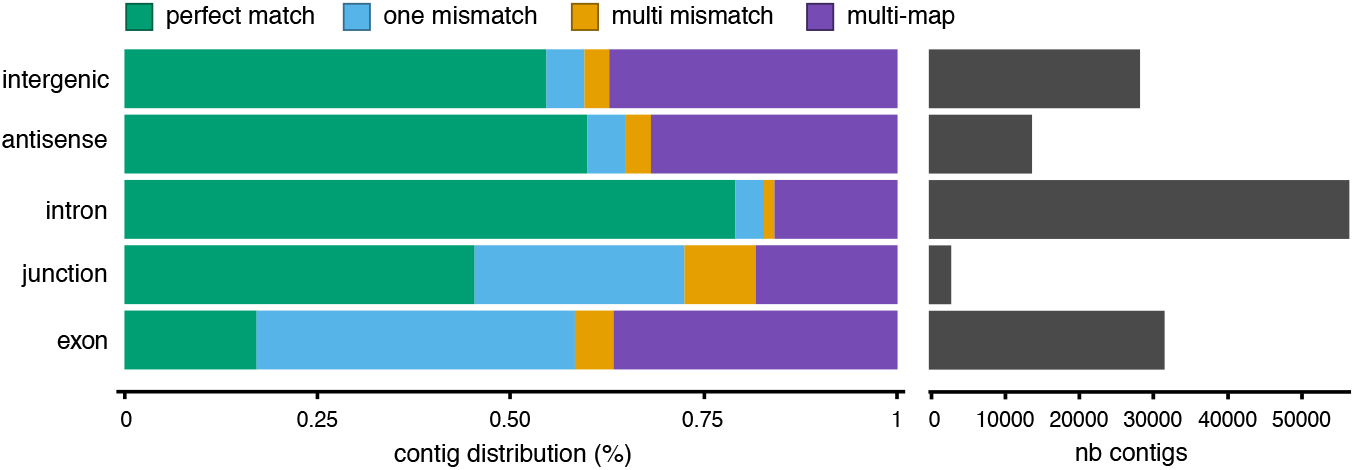
Genomic location of differentially expressed contigs. Contigs are separated by genomic location, according to their overlap with exons, exon-exon junctions, introns, antisense regions of annotated genes or intergenic regions. The right panel shows the total number of contigs in each class; the left panel shows contig distribution according to their alignment status: contigs with a single mapping location are labeled as “perfect match”, “one mismatch” or “multi mismatches”, contigs with multiple mapping locations are labeled as “multi-map”.

### Assigning contigs to biological events

We assigned DE contigs generated from the EMT dataset to 11 classes of potential biological events, using the rule set described in Table 3. Since intragenic DE contigs may result from a mere over/under-expression of their host gene and do not necessarily reflect a differential usage (DU) of transcript isoforms, we implemented a simple strategy to distinguish the two situations based on the differential status of the host gene (Methods). We made this distinction for splicing, polyadenylation, SNVs and intron retention (Table 3).

**Table 3.**
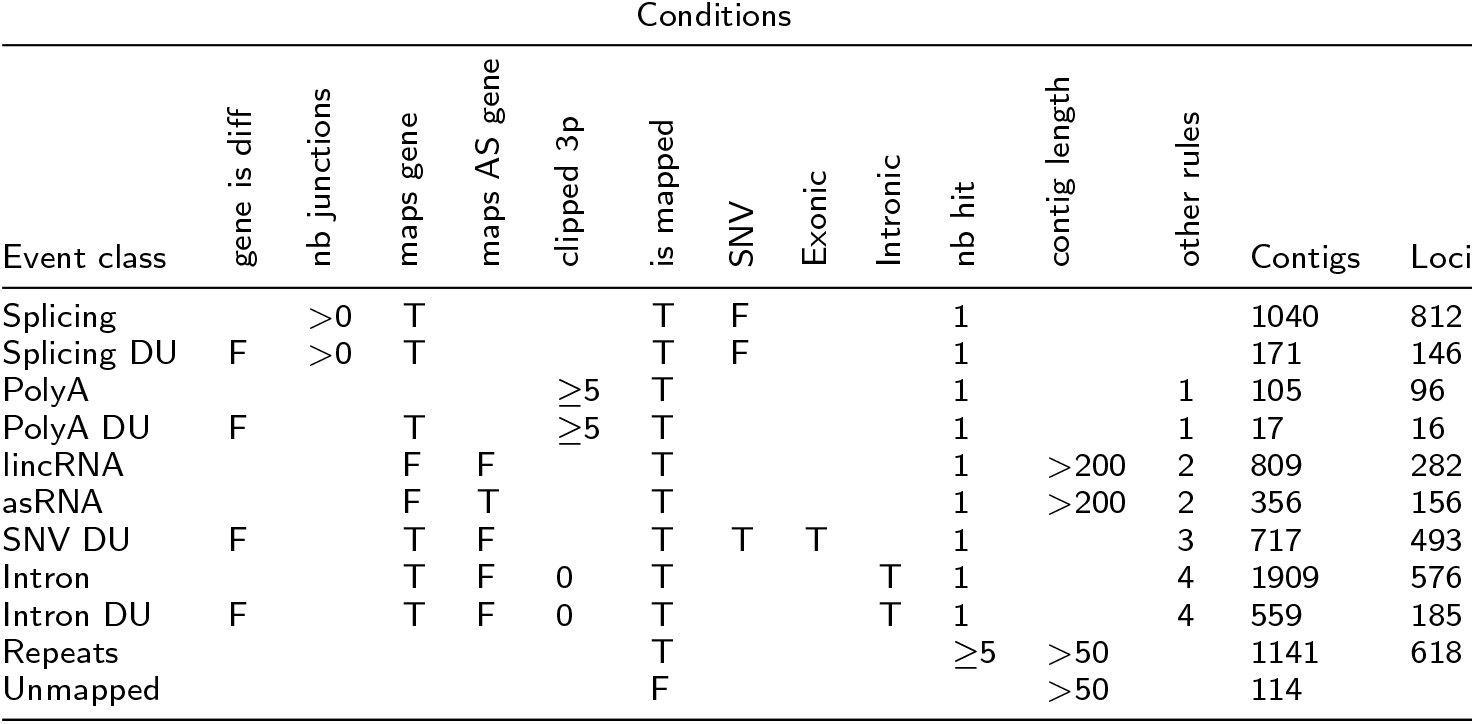
Assignment rules for DE contigs. Each class of event is defined by a set of rules applied to annotated contigs. Keyword “DU” indicates Differential Usage. Column “Other rules” refers to the following: 1. Contig ends with AAAAA, 2. Mean counts in both conditions > 20, 3. Mean counts > 20 in at least 1 condition & mapped region < 1*kb*, 4. Mean counts in cond1+cond2 > 70 & mapped region < 1*kb*. Column “Contigs” indicates the number of contigs of each class found in the EMT experiment. Column “Loci” is the number of loci implicated by these contigs (see Methods).

From the total set of 133k DE contigs (supplemental data), we extracted about 6900 contigs matching our rule set for either event class (Table 3). We noted that a single event often generates multiple contigs. We thus further grouped contigs into “loci”, defined as independent annotated genes or intergenic regions harboring one or more contigs (Table 3). We describe below the main classes of events identified.

#### Differential splicing

Analysis of split-mapped contigs found evidence of potentially novel differential splice variants in 1040 contigs (Table 3, Fig 7A,B,C). Note that this class excludes SNV-containing contigs, as to avoid known splice variants associated to genetic polymorphism. Furthermore, 171 of these contigs were classified as DU, suggesting differential splicing at these sites may not be a consequence from DE of the whole gene. Remarkably, these novel events include a number of subtle variations at 5’ and 3’ splice sites with 3-15 bp difference from the annotated reference, which escaped prior annotation (see eg. Fig S1).

#### Differential polyadenylation

We extracted all contigs aligned with 5 or more clipped (e.g. non-reference) bases at their 3’ end, and containing 5 or more trailing As. Out of 140 such poly-A terminated contigs, 105 (75%) contained an AATAAA or variant polyadenylation signal (Table S1), indicating they result from actual polyadenylated transcripts (Table 3, line “PolyA”). Note these are not necessarily novel polyadenylation sites since polyadenylated transcripts always create k-mers that differ from the reference transcriptome and are hence retained by DE-kupl. Indeed, only 6 of the 105 poly-A contigs mapped intergenic regions. Furthermore, only 17 poly-A contigs mapped to genes with no differential expression (“polyA DU” in 3), and these had relatively poor fold change values (Table S11) raising doubt on their DU status. Altogether this analysis demonstrates that DE-kupl is able to capture *bonafide* polyadenylated transcripts present in the sequencing reads, however we did not observe any clear case of differentially polyadenylated genes in the experiment studied.

#### LincRNA

We identified a subset of 809 DE contigs (282 loci) corresponding to potential long intergenic non-coding RNAs (Table 3, line “lincRNA”). Criteria for lincRNAs were contigs of size > 200nt mapped to an intergenic locus. Visual inspection revealed clear lincRNA-like patterns, whith contigs clustered into well defined transcription units with abundant read coverage and evidence of splicing (Fig 7C, Fig S2). DE-kupl is thus an effective tool for the identification of novel differentially expressed lincRNAs.

#### Antisense RNAs

When DE-kupl is applied to stranded RNA-seq libraries (as with the EMT libraries used in this study), the resulting contigs are strand-specific and can thus be used for identifying antisense RNAs (asRNAs) and disembiguating loci with intricated expression on both strands. We identified 356 contigs from 156 loci mapping to the reverse strand of an annotated gene (Table 3, line “asRNA”). These antisense RNAs include very strong cases of differential expression (Fig 7D), sometimes combined to apparent repression of the sense gene (Fig S3).

#### Allele-specific expression

As DE-kupl quantifies every SNV-containing k-mer, we set out to exploit this capacity to identify potential allele-specific expression events. We extracted all contigs including an SNV (either base substitution or indel) and mapping to an exon whose host gene was not measured as differentially expressed (Table 3, line “SNV DU”). This was a less than perfect procedure, as we did not explicitly test for a switch in allelic balance among the two conditions. Yet, among the 717 contigs identified, some displayed strong apparent changes in allelic balance between the E and M conditions (eg. Fig S4). The ability of DE-kupl to capture differential SNV between datasets may be particularly interesting when looking for recurrent mutations in subpopulations.

#### Intron retention and other intronic events

As highly expressed transcripts often carry intronic byproducts, we expected DE-kupl to turn out a lot of ‘parasitic” intronic contigs. Indeed 1909 contigs mapped to intronic loci (Table 3, line “intron”). We thus focused on intronic k-mers from genes that were not DE (line “intron DU”). This filter identified 559 intronic contigs from 185 different genes. Inspection of read mapping at these loci revealed clear instances of novel skipped of extended exons (Fig S5), as well as cases where a specific short intronic region was differentially expressed, reminiscent of the pattern observed at intronic processed miRNAs and snoRNAs [21] (Fig S6). Therefore DE-kupl can be used for screening a wide variety of exon/intron processing events in addition to alternative splicing.

#### Expressed repeats

Assessing the expression of human repeats by conventional RNA-seq analysis protocols is difficult, as ambiguous alignments render repeat regions “unmappable” [22]. Since DE-kupl first measures expression independently of mapping, we were able to collect and analyze differential contigs with multiple genome hits. 4968 contigs of size 50nt or larger have multiple hits (not shown), and 1141 are repeated more than 5 times (Table 3, line “repeat”). RepeatMasker [23] found 664 out of these 1141 sequences to match known repeats, mostly LINEs, LTRs and SINEs (Fig S8). Further inspection showed that most of the remaining multiple-hit contigs correspond to unannotated repeats or low complexity regions. One of the most striking differential repeats is an unannotated 22x66 bp tandem repeat, located about 2 Mbp from the chromosome 8 telomere. This repeat is found about 50-fold overexpressed in the Mesenchymal condition (Fig 7B, S7). These results indicate DE-kupl can serves as a screen for differential expression or activation of endogenous viral sequences and other repeat-containing transcripts.

**Figure 7.**
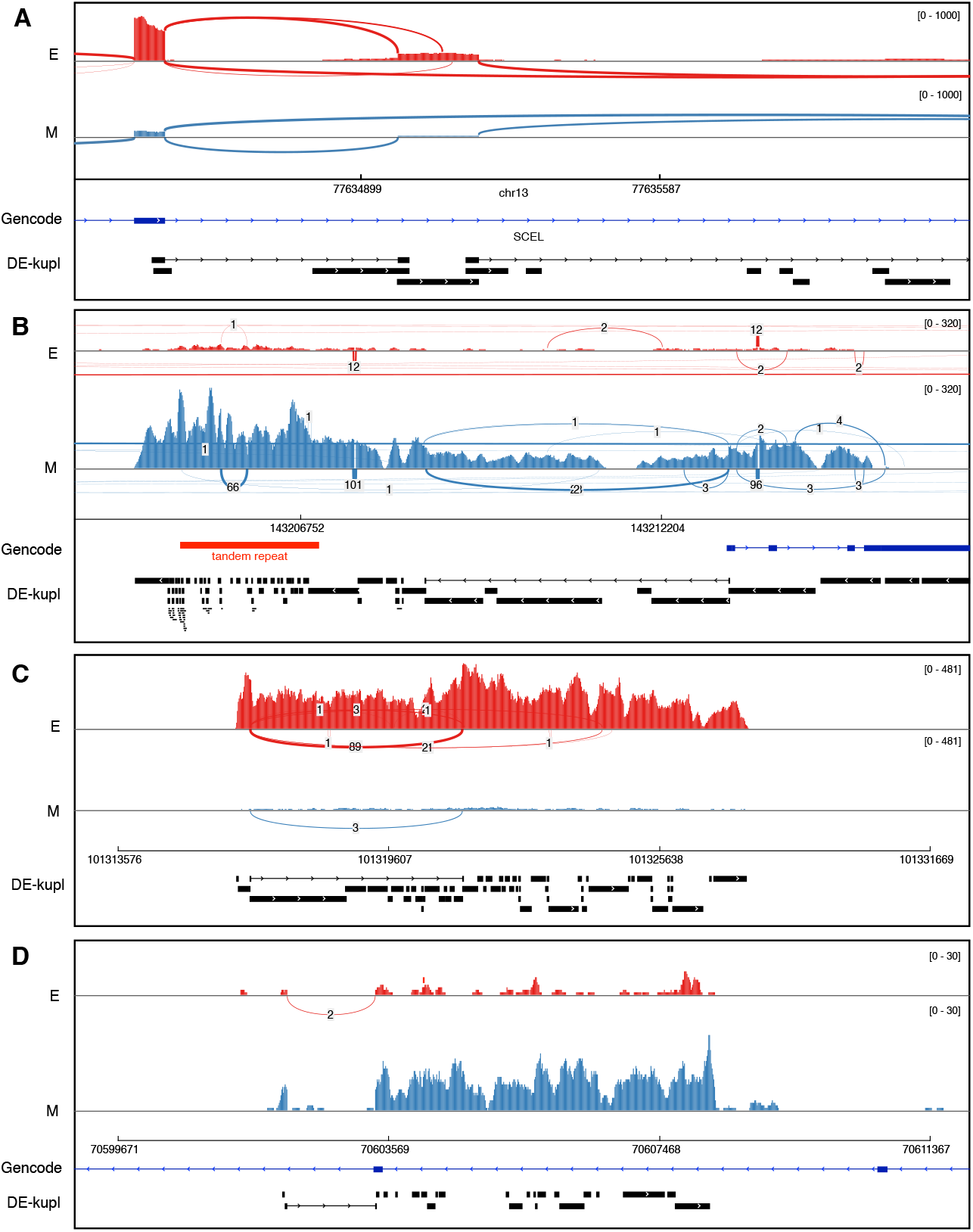
Examples of DE contigs. Sashimi plots generated from IGV using read alignments produced with STAR [36]. Sample SRR2966453 from condition D0 is labeled as “E” (epithelial). Sample SRR2966474 from condition D7 is labeled as “M” (mesenchymal). Annotations from Gencode and DE-kupl DE contigs are shown at the bottom of each frame. **A**. New splicing variant involving an unannotated exon, overexpressed in condition “E”. **B**. Tandem repeat at chr8:143,204-870-143,206,916 (red region) that is overexpressed in condition “M” *vs*. “E”. Note that the overexpressed tandem repeat is part of a larger overexpressed unannotated locus. **C**. A novel lincRNA overexpressed in condition “ E”. **D**. A novel antisense RNA. RNA-seq reads are aligned in the forward orientation while the gene at this locus is in the reverse orientation. The annotated gene is not expressed.

#### Unmapped contigs

Finally, we analyzed DE contigs that did not map the human genome. Unmapped contigs may result from transcripts produced by highly rearranged genes or by exogenous viral genomes and could thus be highly relevant biologically. In principle, DE-kupl is able to detect such events when levels of foreign RNA vary across samples. In this test set, where all samples come from an *invitro* cell line, we did not expect to observe such a phenomenon. Indeed, out of 114 unmapped contigs of size > 50 bp (Table 3, line “Unmapped”), the vast majority (76%) correspond to vector sequences overexpressed in the “M” condition (not shown), indicating these contigs come from the expression vector used for EMT induction. The remaining unmapped contigs were low complexity sequences or aligned to non-human primate sequences, indicating possible contamination and/or misassemblies.

### DE-kupl event detection is reproducible across independent datasets

We cross-validated DE-kupl findings using two independent human RNA-seq datasets extracted from the Genotype-Tissue Expression (GTEx) [24] and the Human Protein Atlas (HPA) [15]. DE contigs were first obtained by running DE-kupl on 8 colon vs 8 skin libraries from GTEx. Events were classified as above into intron retentions, lincRNAs, polyadenylation sites, repeats, splice sites and unmapped. The 100 top events from each class (50 for class ‘unmapped’) were extracted and their k-mer labels saved as a sequence file. We then counted the occurrence of each k-mer in colon and skin libraries from the HPA project and applied DEseq2 [19] to evaluate the significance of the expression change between colon and skin (see Methods). 79% of the 550 DE k-mers identified in GTEX were also significantly DE in the HPA data (Figure 8). Each event class showed clear reproducibility, with particularly strong effects in lincRNAs and splice variants. This demonstrates that novel events identified by DEkupl are reproducible across independent datasets in spite of independent RNA extraction, library preparation and sequencing protocols.

## Discussion

K-mer decomposition followed by filtering and differential expression analysis is a novel way of analysing RNA-seq data that is capable of detecting a wider spectrum of transcript variation than previous protocols. DE-kupl explores all k-mers in the input RNA-seq files (*vs*. only k-mers from annotated transcripts in recent software [8, 9]) which potentially entails heavy computational time and memory requirement. Using the Jellyfish k-mer indexing software and C-programming code for key table manipulation, we achieved time/memory requirements on par with popular mapping-based software for similarly sized datasets. Another key aspect of our protocol that rendered a “full k-mer” analysis tractable was applying successive filters for rare k-mers, Gencode transcripts and differential expression, which altogether resulted in a 200-fold reduction in k-mer counts. These filters are not only useful for technical reasons (they reduce runtimes and enable to get rid of most sequence errors), but they also allow to focus on k-mers which (i) vary significantly between the conditions under study, and (ii) would not be captured by conventional reference-based protocols.

Contrarily to popular RNA-seq analysis software, DE-kupl does not attempt full-length transcript assignment or assembly but focuses instead on local transcript variations. Indeed, we do not consider full-length transcript analysis to be realistic when screening for unspecified RNA variation, since the combinatorial nature of genomic, transcriptomic and post-transcriptomic events would require an indefinitely expanding transcript catalog. In some way DE-kupl is closer in spirit to methods analyzing local RNA-seq coverage such as RNAprof [25] and DERfinder [26], with the notable exception that DE-kupl does not involve mapping and thus avoids mapping-related pitfalls while considerably widening the range of detectable events. Another important benefit of the k-mer strategy is that k-mers representing events of interest can be used to efficiently assess the occurence of similar events in the huge public compendium of RNA-seq data.

We showed that DE-kupl is able to detect a wide range of differential transcription and RNA processing events. Although specialized software may perform better at assessing specific event classes such as differential splicing, to our knowledge no software provides such an extensive screen. As differential RNA-seq analysis is often conducted with an exploratory spirit, we argue that it is preferable to cast a wide net with no preconception on target events, using DE-kupl along with a conventional gene-by-gene differential expression analysis. Note that DE-kupl might also be an interesting option for exploring other types of NGS data such as small-RNA-seq, ChIP-seq or CLIP-seq, with simple adjustments of k-mer size and event annotation rules.

In this proof of concept study, we focused on RNA-seq libraries from a cell line, where no genetic polymorphism was expected among samples. The next step will be application to libraries from multiple individual organisms. Although k-mer diversity will be higher in such datasets, preliminary tests with RNA-seq data from 60 human tumors were completed successfully on a single computer server (data not shown). Analysis of patient samples open exciting perspectives. For instance, the ability of DE-kupl to simultaneously detect genetic variation and RNA expression/processing events may serve as a basis for studying genotype-phenotype relations. Analysis of patient RNA-seq data may also reveal event classes not explored in this work, such as fusion transcripts and circular RNAs.

## Methods

### Characterization of k-mer diversity in human RNA-seq libraries

RNA-seq data for bone marrow, skin and colon were retrieved from the Human Protein Atlas project (10.1126/science.1260419, E-MTAB-2836). A total of 18 datasets from different individuals, corresponding to 6 replicates per tissue, were downloaded from EBI/ENA (bone marrow: ERR315469, ERR315425, ERR315486, ERR315396, ERR315404, ERR315406, colon: ERR315348, ERR315403, ERR315357, ERR315484, ERR315400, ERR315462, skin: ERR315401, ERR315464, ERR315460, ERR315372, ERR315376, ERR315339). The reference GRCh38 genome and Ensembl 86 transcripts were downloaded from Ensembl.

First we counted k-mers in each RNA-Seq and reference sequence set using Jellyfish (2.2.0) count, with options k = 32 and -C (canonical k-mers). The k-mer list for each tissue (Fig 1A and B) was produced by merging counts for all 6 samples and conserving only those found in all replicates.

For mapping statistics (Fig 1B3), we extracted k-mers specific of each tissue and mapped them to the Ensembl 86 transcript reference using Bowtie (version 1.1.2). Unmapped k-mers were mapped a second time with Bowtie to the GRCh38 genome reference. Reads with 3 or more mismatches are not mapped by Bowtie and, therefore, are considered as unmapped.

The intersection of k-mers between RNA-Seq and WGS data (Fig 1C), is based on the transcriptome and genome of lymphoblastoid cell lines [16]. K-mers were counted in these libraries with the same procedure as above. In order to reduce noise from sequencing errors, k-mers with only one occurrence were filtered out.

### DE-kupl Implementation

The DE-kupl pipeline (Fig S12) is implemented using the Snakemake [27] workflow manager. A configuration file is filled up by the user with location of FASTQ files, the condition of each sample, as well as global parameters such as k-mer length, CPU number, maximum memory and other parameters for each step of the pipeline, as described hereinafter.

#### K-mer counting

Raw sequences (FASTQ files) are first processed with the `jellyfish count` command of the Jellyfish software, which produces one index (a disk representation of the Jellyfish hash-table) for each sequence library. For stranded RNA-seq libraries, reads in reverse direction relative to the transcript are reverse-complemented, ensuring proper orientation of k-mers. At this point, for each library, only k-mers having at least 2 occurrences are recorded (user-defined parameter). Once a Jellyfish index is built, we use the `jellyfish dump` command to output the raw-counts in a two column text file, which contains at each line a k-mer and its number of occurrences. Raw counts are then sorted alphabetically by k-mer sequence with the Unix sort command.

#### K-mer filtering

All samples counts are joined together using the `dekupl-joinCounts` binary to produce a single matrix will all k-mers and their abundance in all samples. Given an integer *a* ≥ 0, we define the *recurrence* of a *k*-mer *x* as the number of samples where *x* appears more than *a* times, i.e. *recurrence* 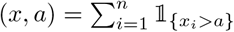, where *n* is the total number of samples and *x_i_* is the number of times the *k*-mer *x* appears in sample *i*. The *k*-mer filtering step involves two user-defined parameters: an integer `min_recurrence_abundance` and an integer `min_recurrence` such that a *k*-mer *x* is filtered out if *recurrence*(*x*, `min_recurrence_abundance`) < `min_recurrence`, i.e. if the *k*-mer*x* appears more than `min_recurrence_abundance` times in fewer than `min_recurrence` of the samples. Usually `min_xecurrence` is set to the number of replicates in each conditions, and `min_recurrence_abundance` is set to 5. In order to remove known transcripts sequences from our set of experimental k-mers, we also use our Jellyfish-based procedure to create the set of *k*-mers appearing in the reference transcriptome and we subtract this set from the experimental k-mers.

#### Differential k-mer expression

Prior to differential analysis, we compute normalization factors (NFs) using the “median ratio method” [28] on the table of k-mers after recurrence filter: for each sample, the NF is the median of the ratios between sample counts and counts of a pseudo-reference obtained by taking the geometric mean of each k-mer across all samples. To avoid dealing with the complete table of k-mers, we extracted a random subset of 30% of the k-mers and computed NFs on this subset. Computing NFs on the complete table of k-mers, on the table of k-mers after recurrence and Gencode filters, or on the table of transcripts abundances produced by Kallisto [8] led to similar values (Fig S9).

To perform differential analysis, two options are implemented (Fig. S10). The first option is to apply a t-test for each k-mer on the log transformed counts, normalized with the previously computed NF. Transformation of raw counts in conjunction with linear model analysis have been successfully used for differential analysis of counts [29]. We perform the t-test independently on each k-mer and avoid complex variance modeling strategies to reduce execution time of the analysis. The t-test option has been implemented in C in the `dekupl-TtestFilter` binary. Note that this *t*-test option is not appropriate for small samples [30]. To increase the power of the analysis, in particular for small samples (typically less than 6 vs 6 libraries), we strongly advise to use the second option based on generalized linear model, implemented in the `R` package `DESeq2` [19]. On top of modeling raw counts (normalization or prior log-transformation of the counts is not required), this approach performs information sharing across k-mers to improve variance estimation and differential analysis results. However, given the large number of k-mers, we do not apply this approach on the complete matrix of k-mer counts. We divide the matrix of k-mer counts into random chunks of approximately equal size (around one million k-mers) and apply the DESeq2 approach on each chunk independently. For each chunk, previously computed NFs are used as an input of the method, and are not computed independently on each chunk. Raw p-values, not adjusted for multiple testing, are collected as an output for each chunk, and merged into one single vector containing the raw p-values for all k-mers to test. Subsequently, raw p-values obtained from either the *t*-test or the `DESeq2` test are adjusted for multiple comparisons using the Benjamini-Hochberg procedure [31] and k-mers with adjusted p-values above a user-set cutoff are filtered out.

#### K-mer assembly

DE k-mers are assembled *de novo* in order to group k-mers that potentially overlap the same event (ie. all k-mers overlapping a splice junction or SNV). To this aim, we developed our own procedure called `mergeTags`, which works as follows: we first identify all exact *k*−1 prefix-suffix overlaps between k-mers. We consider only k-mers that overlap with exactly one other k-mer, and merge all pairs of k-mers involved in such overlaps. For example, given the set of k-mers: *ATG*, *TGA*, *TGC*, *CAT*, the following contigs are produced: *contigs* = *CATG*, *TGA*, *TGC*. We repeatedly merge contigs that overlap exactly over *k* − 1 bp with exactly one other contig. We then repeat this assembly process with *k* − 2 exact prefix-suffix overlaps, using as input the contigs produced at the previous step, and so forth for increasing values of *i* such that *k* − *i* > 15 bp (default value). Finally, a set of DE contigs is produced and each contig is labelled by its constitutive k-mer of lowest p-value. This assembly procedure is implemented in C in the `dekupl-mergeTags binary`.

#### Contig Annotation

Finally, DE contigs are annotated in order to facilitate biological event identification. First, contigs are aligned with BLAST [32] against Illumina adapters. Contigs matching adapters are discarded. Retained contigs are further mapped to the reference Hg38 human genome using the GSNAP short read aligner [33], which showed the best speed/sensitivity ratio for aligning both short and long contigs in internal tests (not shown). GSNAP is used with option `-N 1` to enable identification of new splice junctions. Contigs not mapped by GSNAP are collected and re-aligned using BLAST.

Alignment characteristics are extracted from GSNAP and BLAST outputs. Alignment coordinates are compared with Ensembl (v86) annotations (in GFF3 format) using BEDTools [34] and a set of locus-related features is extracted. The final set of annotated features (Table S3) is reported in a contig summary table. The annotation procedure generates two additional files: a “per locus” summary of contigs (one line per genic or intergenic locus), and a BED file of contig locations that can be used as a display track in genome browsers. In the “per locus” table, a locus is defined as either an annotated gene, the genomic region located on the opposite strand of an annotated gene, or the genomic region separating two annotated genes. The table records the number of contigs overlapping each locus as well as the contig with lowest FDR for this genomic interval.

In parallel to k-mer counting and filtering, we analyze the RNA-Seq data libraries a conventional differential expression protocol. Reads are processed with Kallisto [8] to estimate transcript abundances. Transcript-level counts are then collapsed to the gene-level and processed with DESeq2 [19] to produce a set of differentially expressed genes. This information is stored in the contig summary table and used later on for defining events with differential usage (“DU” in Table 3).

#### DE-kupl run on EMT data

DE-kupl was run using RNA-seq libraries from [20] retrieved on the GEO web site under accession GSE75492. For stage “E” we used libraries GSM1956974, GSM1956975, GSM1956976, GSM1956977, GSM1956978, GSM1956979, and for stage “M” GSM1956992, GSM1956993, GSM1956994, GSM1956995, GSM1956996, GSM1956997. DE-kupl parameters were `kmer_length 31, min_recurrence 6, min_recurrence_abundance 5, pvalue_threshold 0.05, lib_type stranded, diff_method Ttest`. Output files are provided as supplementary material. The DE-kupl contig summary table was analyzed interactively using R commands to extract lists of contigs based on the filtering rules described in Table 3. Visualization of selected contigs was performed with IGV [35], using the bed file produced by DE-kupl and read mapping files produced by STAR [36].

#### Cross-validation

DEkupl was applied to 8 skin and 8 colon libraries from GTEx [24]: skin library IDs: SRR1308800, SRR1309051, SRR1309767, SRR1310075, SRR1311040, SRR1351501, SRR1400467, SRR1479595; colon library IDs: SRR1316343, SRR1396146, SRR1397292, SRR1477732, SRR1488307, SRR807751, SRR812697, SRR819486. DE-kupl parameters were `kmer_length 31, min_recurrence 6, min_recurrence_abundance 5, pvalue_threshold 0.05, lib_type unstranded, diff_method Ttest`. DE-kupl contigs were interactively classified using R commands, applying the same rules as in Table 3. Classes asRNA and SNV-DU were not included since asRNA identification is not possible using the unstranded GTEx and HPA libraries, and we had no reason to expect common SNVs with differential usage in this dataset. DE contigs were sorted by fold-change and k-mer labels of the top 100 DE contigs in each class were extracted (50 for class ‘unmapped’ due to lower event number). DE-kupl output files and selected k-mers are provided in supplementary material. For cross-validation, we used the same 6 skin and 6 colon RNA-Seq data as in Figure 1 (10.1126/science.1260419, E-MTAB-2836). K-mers were counted in each library using Jellyfish with options `k = 31` and `-C` (canonical k-mers) as GTEx data were unstranded. All k-mers selected from the GTEx analysis were queried against the jellyfish databases using `jellyfish query` command. Finally the extracted k-mers counts were processed with DESeq2 [19] and the resulting adjusted p-values were plotted for each event class (Figure 8).

**Figure 8.**
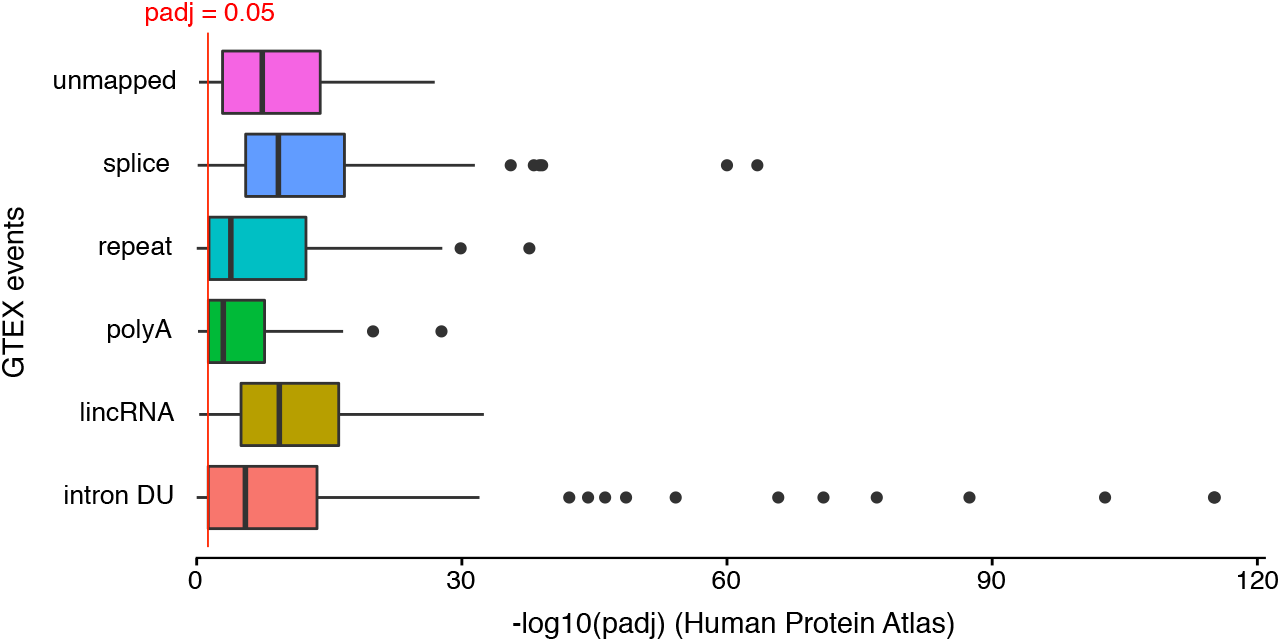
Cross validation of DE events in GTEx and Human Protein Atlas data. 550 DE contigs from 6 different event classes (intron with differential usage, lincRNA, polyA site, repeat, splice site, unmapped) were identified using DE-kupl on GTEx libraries from two human tissues (skin and colon). A representative k-mer from each contig was then tested for differential expression in skin and colon libraries in the Human Protein Atlas. Box plots represent distributions of DESeq2 adjusted P-values for all k-mers in the different classes. The red line shows the adjusted P-value cutoff of 0.05.

#### Availability

The DE-kupl software, documentation and supplemental material presented herein are available at https://transipedia.github.io/dekupl/.

## Competing interests

The authors declare that they have no competing interests.

## Author’s contributions

JA, NP, RC, MS, MeG, TC and DG designed the study. JA, MaG, MeG and JLC developed the code. JA, NP, RC, MS, MeG, TC and DG analyzed the results. DG and JA drafted the manuscript and all authors proofread and corrected the manuscript.

## Acknowledgements

We thank Damien Drubay for useful statistical discussions and Emilie Drouineau for setting up the DE-kupl Docker container. This project was supported in part by grants “Plan Cancer – Systems Biology” #bio2014-04 and ANR-10-INBS-0009 (France Génomique) from Agence Nationale pour la Recherche to DG. GTEx data was downloaded from the dbGaP web site under phs000178/GRU.

